# MC profiling: a novel approach to analyze DNA methylation heterogeneity from bulk bisulfite sequencing data

**DOI:** 10.1101/2022.07.06.498979

**Authors:** Giulia De Riso, Antonella Sarnataro, Giovanni Scala, Mariella Cuomo, Rosa Della Monica, Stefano Amente, Lorenzo Chiariotti, Gennaro Miele, Sergio Cocozza

## Abstract

DNA methylation is an epigenetic mark implicated in crucial biological processes. Most of the knowledge about DNA methylation is based on bulk experiments, in which DNA methylation of genomic regions is reported as average methylation. However, average methylation does not inform on how methylated cytosines are distributed in each single DNA molecule.

Here, we propose Methylation Class (MC) profiling as a genome-wide approach to the study of DNA methylation heterogeneity from bulk bisulfite sequencing experiments. The proposed approach is built on the concept of MCs, groups of DNA molecules sharing the same number of methylated cytosines. The relative abundances of MCs from sequencing reads incorporates the information on the average methylation, and directly informs on the methylation level of each molecule.

By applying our approach to publicly available bisulfite-sequencing datasets, we individuated cell-to-cell differences as the prevalent contributor to methylation heterogeneity. Moreover, we individuated signatures of loci undergoing imprinting and X-inactivation, and highlighted differences between the two processes. When applying MC profiling to compare different conditions, we identified methylation changes occurring in regions with almost constant average methylation.

Altogether, our results indicate that MC profiling can provide useful insights on the epigenetic status and its evolution at multiple genomic regions.

## INTRODUCTION

DNA methylation is a heritable epigenetic mark consisting of the enzyme-mediated addition of a methyl-group to deoxyribonucleotides (1–3). In mammals, DNA methylation mainly involves cytosines in CpG context (1–3). DNA methylation has been shown to regulate gene expression and genome stability, and has been implicated in crucial biological processes, like genomic imprinting and X inactivation (3–6). In cell differentiation, DNA methylation shapes fate and engraves the identity of cells (1,7). Its dysregulation has been linked to plenty of pathological conditions (8–11).

Several experimental techniques have been developed to study DNA methylation (12). Among them, bisulfite sequencing techniques are widely adopted to assess the methylation status at single base resolution, either at targeted regions or at genome-wide level (12–16).

Single base DNA methylation is usually reported as the fraction of molecules in which a given cytosine is methylated (17). Genome-wide methylation analysis has highlighted that most cytosines are not evenly methylated in different molecules (18). Cellular heterogeneity and allele specific methylation are potential sources of this molecular heterogeneity (19).

Evidence has been provided that DNA methylation is regulated in larger genomic regions, with sets of neighboring cytosines working as functional units (20–23). DNA methylation analysis has indeed turned to the study of the average methylation of DNA regions (the fraction of methylated cytosines in a given region), and on the identification of regions with consistently different DNA methylation levels between groups of samples (differentially methylated regions, DMR) (17). This latter approach has largely contributed to the current knowledge on DNA methylation and its implication in health and disease status (24–26).

However, the overall average methylation of a region does not inform on how this amount is contributed by the single DNA molecules. As an example, an average methylation value of 0.5 for a given locus could result from a homogenous pool of half methylated molecules, or from an heterogeneous, balanced set composed of fully methylated and unmethylated molecules, or even from more heterogeneous pools (Figure 1 of (27)).

**Figure 1:**
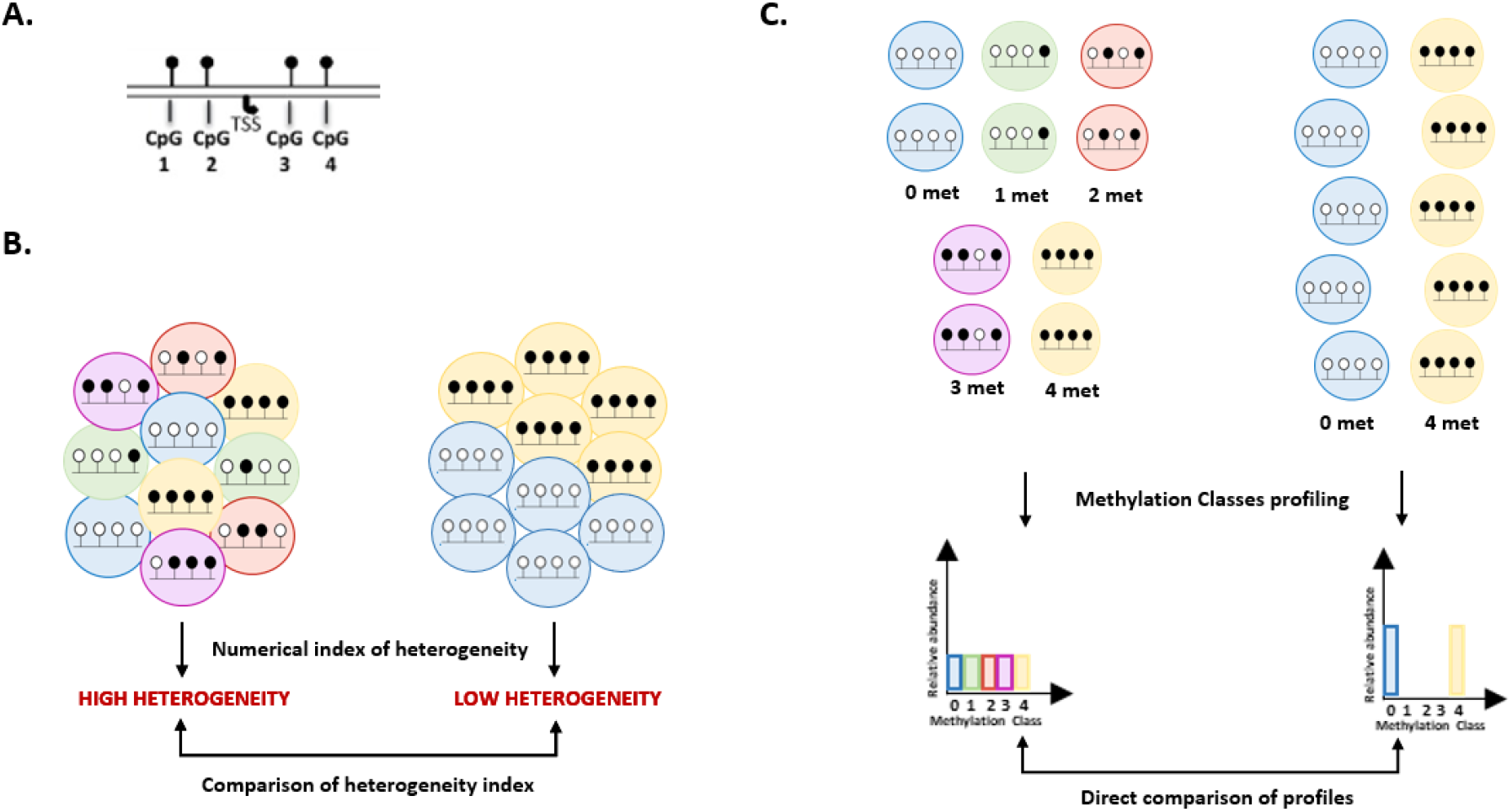
rationale of MC profiling. **A**: example of a region of interest holding 4 CpGs (TSS=Transcription Starting Site) **B**: Representation of epiallele-based analysis. The DNA methylation heterogeneity for a certain locus is usually quantified through a numerical index (e.g., epipolymorphysm, Shannon entropy, ‥). This can be then adopted to compare the heterogeneity of pools of molecules (for example, to compare the heterogeneity of a certain locus in different samples). **C**: Representation of MC profiling analysis. The epialleles are first grouped in Methylation Classes (MCs) according to the number of methylated cytosines. The relative abundances of the possible MCs (for a locus holding n CpGs there are n+1 possible MCs), named MC profile, summarize the molecular heterogeneity and the methylation levels of a given region. MC profiles can be directly adopted to perform differential analysis.

Single-cell DNA methylation assays have highlighted extensive cell-to-cell differences in regional DNA methylation (28,29), and have demonstrated that cellular heterogeneity can have a functional impact. For example, epigenetic variability at regulatory elements has been linked with gene expression variability (30,31). However, single-cell DNA methylation assays are still limitedly adopted due to the high cost and large sparsity of produced data (28,32).

Besides single cell techniques, analysis of DNA methylation patterns in bisulfite sequencing reads has also been adopted to analyze DNA methylation heterogeneity in bulk samples (19,28,33–36). In fact, analyzing the arrangements of methylated and unmethylated cytosines (usually referred to as epialleles) in a DNA molecule can provide information on cellular heterogeneity of a sample. Several scores have been developed to quantify this cellular heterogeneity and perform differential analysis of epiallele composition among conditions (19). In this context, the different epialleles are usually regarded as independent elements, and the overall amount of methylated cytosines (i.e., methylation level) is not, or only partially, taken into account (33–37).

It could be reasonable to hypothesize, however, that the functional properties of the different epialleles can be at least affected by their methylation level, and that epialleles with the same amount of methylated cytosines (regardless of their position) share similar functional properties. Therefore, examining the distribution of groups of epialleles that share the same methylation level (Methylation Classes MCs), rather than of epialleles themselves, can add functional information to the study of the epigenetic heterogeneity of a sample.

Following this approach, mathematical modeling has been applied to estimate the distribution of single-molecule methylation levels from Whole Genome Bisulfite Sequencing (WGBS) data, giving insights on its disposition across the genome, its evolution upon differentiation, aging and cancer, and its relationship with the genetic background (38–40).

In previous studies, we directly estimated the distribution of MCs at targeted loci analyzed through high-depth amplicon bisulfite sequencing, thus obtaining information on the regulatory mechanisms of DNA methylation (41,42).

In this study, we extended the concept of MCs to genome-wide bisulfite sequencing data, and developed a method named MC profiling with the aim to explore DNA methylation heterogeneity at multiple target regions.

MC profiling identified cell-to-cell differences as the prevalent contributor to DNA methylation heterogeneity, with allele differences emerging in a small fraction of analyzed regions. Moreover, MC profiling led to the identification of signatures of loci undergoing genomic imprinting and X inactivation, and highlighted differences between the two processes. When applied to a dynamic system, MC profiling identified DNA methylation changes in regions with almost constant average methylation. Altogether, our results indicate that MC profiling can provide useful insights on the epigenetic status and its evolution at multiple genomic regions.

## RESULTS

### The MC profiling approach

#### a. Rationale of MC profiling

The rationale of MC profiling, and the differences with epiallele-based approaches, is depicted in Figure 1.

Epiallele-based approaches are based on the analysis of the arrangements of methylated and unmethylated cytosines (epialleles) in sequencing reads mapping to a region of interest, like the one depicted in Figure 1A. Considering each reads coming from a DNA molecule, several scores have been developed to quantify the heterogeneity observed in a bulk sample, and to compare it among different samples (Figure 1B) (19). This approach has proved to be particularly suitable, for example, to individuate regions undergoing clonal selection and epigenetic drift in tumors (33,34,37). However, the methylation level of epialleles is usually not, or only partially, incorporated in these heterogeneity scores (33–37).

The underlying idea of our approach is, instead, that looking at the distribution of epialleles grouped by their methylation levels adds useful information for the functional interpretation of DNA methylation heterogeneity in a sample.

MC profiling is based on the empirical estimate of the distribution of epialleles grouped by their methylation levels (Figure 1C). MC profiling uses data from Reduced Representation Bisulfite Sequencing (RRBS) experiments, thus allowing for the simultaneous analysis of thousands of regions from the same sample. Instead of adopting a numerical index (as, for example,the Shannon Index) to summarize the DNA methylation heterogeneity of a given region, we kept as much information as possible and described, for each DNA region, the relative abundance of the possible MCs. In this setting, we adopted the direct comparison of MC profiles to analyze differential methylation of a given region among conditions, or to examine the differences among regions in the same condition (Figure 1C). Comparing MC profiles allowed us to compare not only the heterogeneity but also the different methylation levels of DNA molecules.

In summary, adopting MC profiles can provide the following advantages:

– Considering how they are computed, MC profiles directly incorporate the average methylation of a given region, and inform on how it is contributed by single DNA molecules.
– MC profiles retain all information from a pool of molecules, and enable the direct visualization of DNA methylation heterogeneity of a given region
– MC profiles are empirically estimated from sequencing reads, and are independent on a priori parametrization of DNA methylation dynamics (see Discussion

#### b. Description of MC profiling

In this study, we adopted MC profiles to represent the DNA methylation status of a region of interest. In particular, we focused on non-overlapping regions made up of 4 CpG sites, moving from the observation that the number of reads per region drops when increasing the number of CpG sites from 4 to 5 (34). We will refer to these regions as *epiloci* in the following text (Figure 2A). Based on the results obtained on simulated data (see Methods), we selected for MC profiling those epiloci covered by at least 50 reads, considering only reads spanning the entire epilocus.

**Figure 2:**
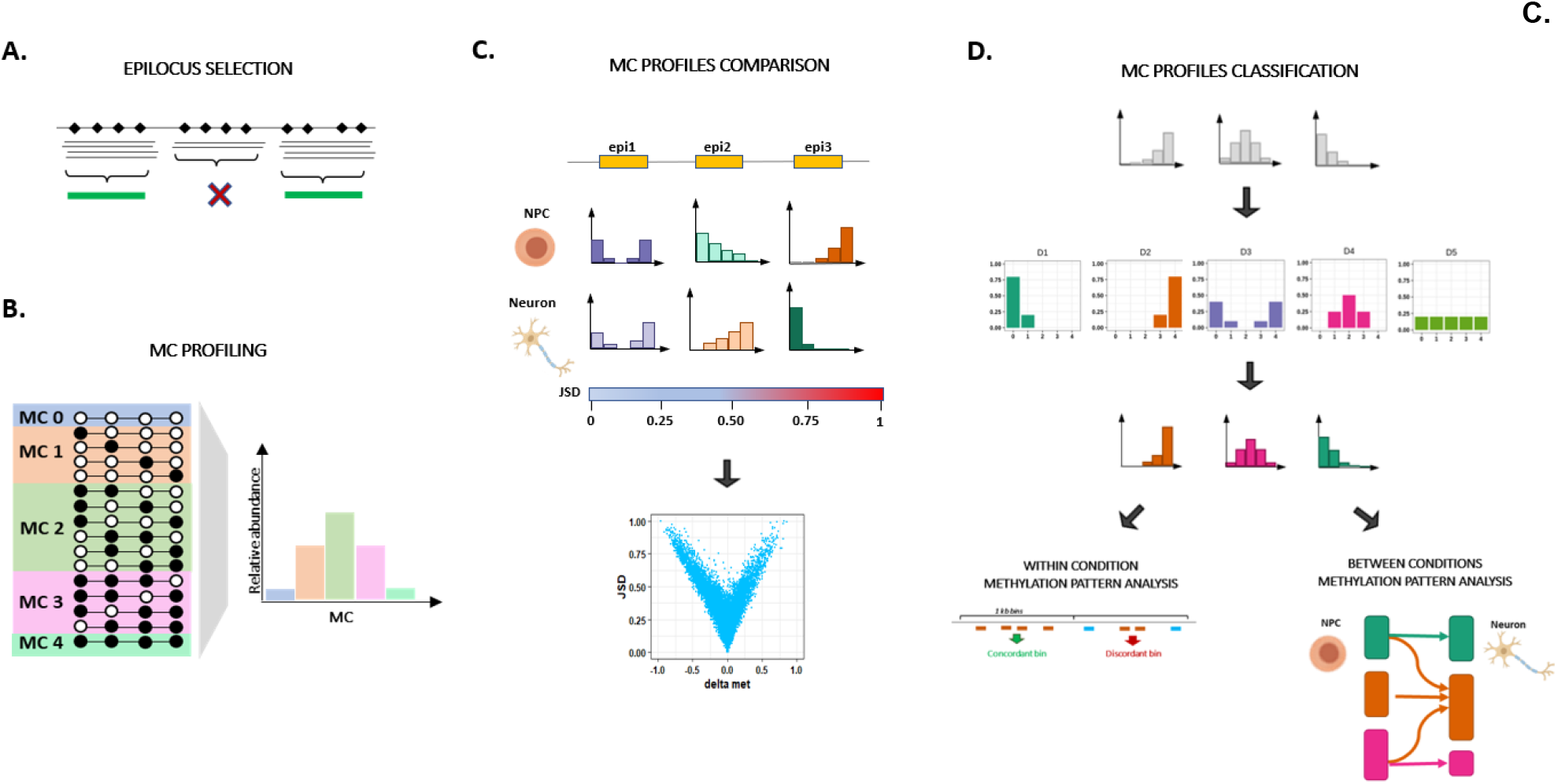
schematic drawing of MC profiling. **A**: Selection of epiloci eligible for MC profiling. An epilocus is defined as a genomic region holding 4 CpGs. Non-overlapping epiloci with coverage higher than 50 reads were retained for MC profiling. **B**. For a given epilocus, the MC profile is computed as the fraction of reads supporting the possible MCs out of the total number of reads. Only reads spanning the entire epilocus were considered in the computation of the MC profile. **C**. The Jensen-Shannon distance is used to quantify the degree of dissimilarity between MC profiles. Based on the results obtained from simulated data, we considered two MC profiles to be different when observing a JSD above 0.26. The JSD can be used to assess the changes of MC profiles at a given epilocus in different conditions. The JSD can be also compared to other metrics, such as the difference of average methylation (delta met), over the analyzed epiloci. **D**. MC profiles were assigned to 5 Methylation Patterns (MPs) according to the most similar among 5 archetypal profiles (here indicated in the upper panel, middle row). This data compression procedure provided us with a signature of genome-wide MC profiles composition in a given condition. MPs enabled us to i) directly compare the MP of different epiloci within the same sample/ or condition (within sample analysis) and ii) to compare the MP transitions occurring at a given epilocus in different conditions (between conditions analysis).

For a certain epilocus, methylated and unmethylated cytosines can be arranged in 16 possible combinations, named epialleles, in a single DNA molecule (Figure 2B). Following the hypothesis that molecules with the same amount of DNA methylation may share functional properties, we grouped epialleles into 5 methylation classes (MCs) according to the number of methylated cytosines they bear. Hence, for a given epilocus, we obtained the MC profile from the relative abundance of the possible MCs (Figure 2B).

To quantify the dissimilarity between two MC profiles, we adopted the Jensen Shannon Distance (JSD, see Methods). Based on the results obtained on simulated data (see Methods), we considered two MC profiles to be different when we observed a JSD above 0.26. Through JSD we could, for example, compare the MC profiles of a given epilocus in different conditions. We could also compare the JSD with the overall average methylation over the analyzed epiloci (Figure 2C).

To further improve the interpretability of the data, we adopted a data compression procedure, and assigned each MC profile to a Methylation Pattern (MP) according to the most similar of 5 archetypal profiles, hereafter referred to as prototypes (Figure 2D, see Methods). The prototypes, which are reminiscent of standard discrete distributions, were chosen because they reflect the reasonable profiles of an epilocus expected at a given methylation amount. In fact, D1 and D2 represent the two symmetric profiles for highly methylated or unmethylated epiloci, in which we expect a prevalence of fully unmethylated and methylated MCs, respectively. D3, D4 and D5, instead, represent the hypothetical profiles of intermediately methylated regions, that can reflect 1) the prevalence of both fully methylated and unmethylated MCs (D3, bimodal profile), 2) the prevalence of intermediately methylated MCs (bell-shaped profile, D4), or 3) the presence of all possible MCs with the same relative abundance (uniform profile, D5).

The data compression procedure provided us with a signature of MC profiles composition over all the analyzed epiloci of a sample. Importantly, this signature did not depend on the specific experimental system. In this way, we were able to i) directly compare the MP of genome-wide epiloci within the same sample/condition and ii) to compare the MP transitions occurring at a given epilocus in different conditions (Figure 2D).

#### c. MC profiles conjugate quantitative methylation and molecular heterogeneity of an epilocus

We applied MC profiling to 2 datasets of samples publicly available in GEO (see Materials and Supplementary Table 1). Dataset1 included samples from 3 wild-type mice embryos, whereas Dataset2 included 3 samples from human CD19+ B-cells isolated from normal controls. Indeed, our datasets came from different species and were representative of different developmental stages, where we expect that DNA methylation heterogeneity probably derives from different dynamics (epigenetic drift in somatic cells vs cell differentiation in mouse embryos). We reasoned that such an experimental plan would have enabled us to generalize the results of our analysis.

For each sample, we profiled about 100000 epiloci in Dataset 1 and 90000 epiloci in Dataset 2. The systematic description of the analyzed cytosines is reported in Supplementary Figure 1-2.

By examining the average methylation of epiloci belonging to different MPs, we confirmed that the quantitative amount of methylated cytosines of assigned elements was coherent with the expected values for each pattern (Figure 3A, Supplementary Figure 3). However, MC profiles add further information depicting the heterogeneity of DNA methylation among DNA molecules. This was particularly evident for the D3, D4, and D5 patterns. In fact, epiloci exhibiting the same average methylation were assigned to different MPs (Figure 3B).

**Figure 3:**
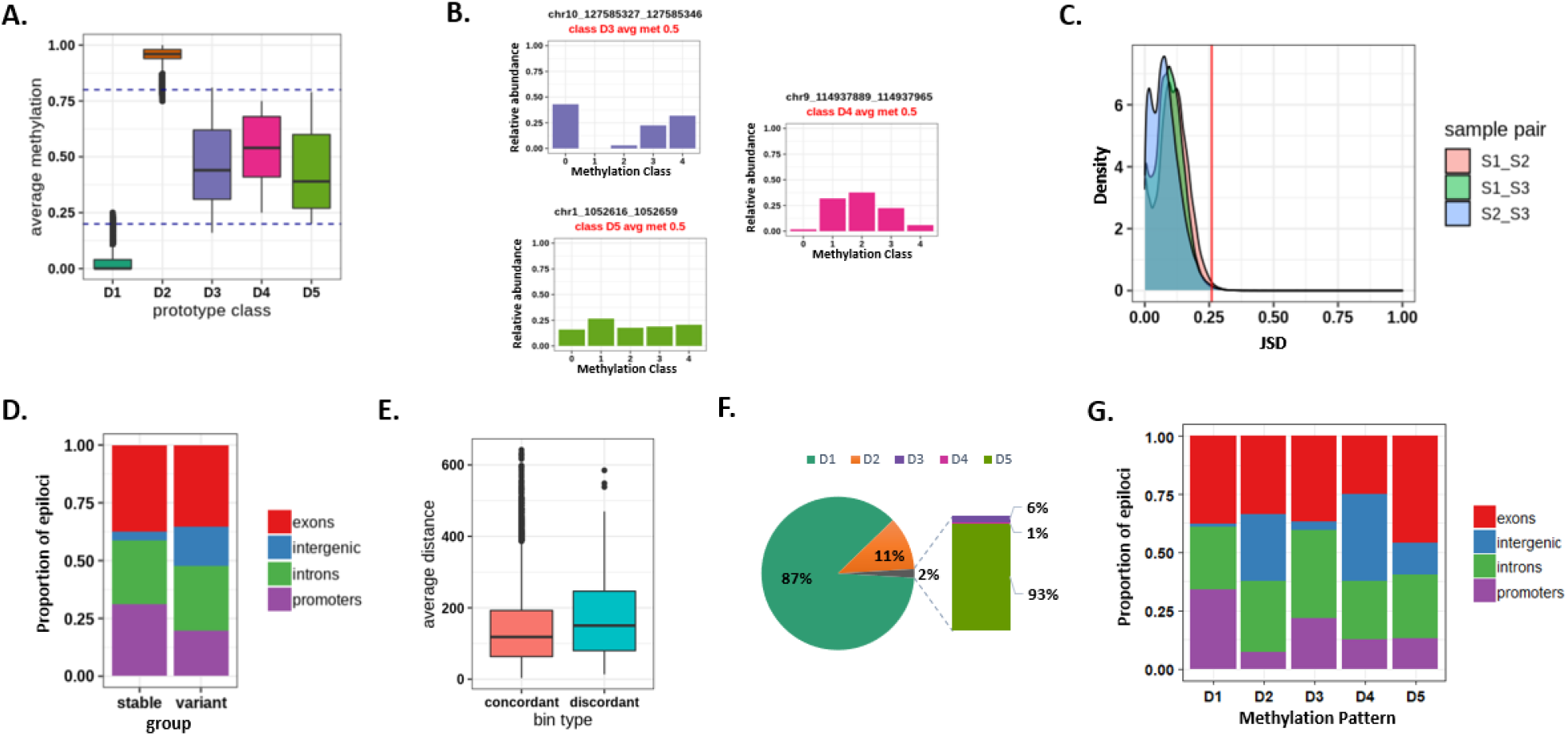
MC profiling results for Dataset 1. **A**: Average methylation level of epiloci assigned to each MP. **B**: example of epiloci with same average methylation and different MC profiles. **C**: Density plot of MC profile distance between sample pairs. x-axis: JSD values between sample pairs. y-axis:density of epiloci with a given sample-pairs JSD value The red line indicates the cutoff value of JSD. **D**: Genomic annotation of epiloci with stable or variant MC profiles. **E**: average distance between epiloci inside concordant and discordant bins. **F**: Fraction of epiloci attributed to the different MPs. **G**: genomic annotation of epiloci assigned to the different MPs.

#### d. MC profiles are mostly stable among individuals and across genomic regions

We investigated the stability of MC profiles across samples. For this aim, we analyzed the epiloci for which the MC profiles were assessed in all the samples in the individual datasets (n=87457 and n=41609) and computed the JSD of MC profiles among sample pairs. We found that 98% of epiloci in Dataset 1 and 96% in Dataset 2 had JSD lower or equal to 0.26 in all sample pairs, meaning that MC profiles at most of the epiloci were very similar between samples (Figure 3C, Supplementary Figure 4A).

In both datasets, we found that stable epiloci, i.e. epiloci with a JSD below the cutoff in all sample pairs (n=86319 and n=39767, in Dataset1 and 2 respectively), were enriched in promoters (chi-square post hoc test p-values < 1e-7) and depleted in intergenic regions (chi-square post hoc test p-values < 1e-7). On the contrary, variant epiloci, i.e. epiloci with JSD above the cutoff in at least one sample pair (n= 1138 and n=1842, in Dataset 1 and 2 respectively), were depleted in promoters (chi-square post hoc test p-values < 1e-7) and were enriched in intergenic regions (chi-square post hoc test p-values < 1e-7). We found no difference in the proportion of stable and variant epiloci located in coding sequences (Figure 3D, Supplementary Figure 4B).

Based on this result, for each dataset we retained for further analysis the stable epiloci, and computed the consensus MC profile by averaging the relative abundances of each MC from the three samples. We then applied the data compression procedure, and assigned the consensus MC profiles to the MPs (see Methods).

Since epigenetic modifications are expected to involve larger DNA regions than individual epiloci, we expected that neighboring epiloci exhibited concordant MC profiles. To test this hypothesis, we binned the genome in 1 kb regions, and compared the MPs of epiloci located in each bin (see Methods). Among the bins harboring at least 3 epiloci, 10733 (99%) bore concordant and (1%) 148 bore discordant epiloci in Dataset 1, whereas (95%) 5079 bore concordant and (5%) 249 bore discordant epiloci Dataset 2. We confirmed that the number of bins bearing concordant epiloci significantly differed from the one expected by chance in both datasets (see Methods and Supplementary Figure 5). This result suggests that MC profiles of neighboring epiloci tend to be similar. This conclusion is further supported by the observation that the reciprocal distance between epiloci in concordant bins tends to be lower than in discordant bins (Mann-Whitney p-value = 0.0005866, Figure 3E, Supplementary Figure 4C).

Overall, MC profiles resulted to be mostly stable among individuals and across genomic regions, thus suggesting that the heterogeneity captured by MC profiles mostly results from controlled DNA methylation dynamics, rather than from stochastic fluctuations of methylation levels.

#### e. MC profiles differentiate functional genomic regions

We reasoned that assigning MC profiles to different MPs could provide us with a signature of genome-wide MC profiles composition in a given dataset. We indeed examined the proportion of epiloci assigned to each MPs. In accordance with the well-established bimodal distribution of average DNA methylation (16), the most represented prototype classes were D1 (83% and 78% of epiloci in Dataset 1 and 2, respectively) and D2 (about 15% and 16% of epiloci in Dataset 1 and 2, respectively). The intermediately methylated D3, D4 and D5 classes accounted respectively for 2% of epiloci in Dataset 1 and 5% of epiloci in Dataset 2 (Figure 3F, Supplementary Figure 4D). Among the intermediately methylated classes, the most represented one was the D5 (90% and 82% of epiloci in Dataset 1 and 2, respectively), followed by the D3 class (9% and 13% of epiloci in Dataset 1 and 2, respectively). The D4 class was strongly underrepresented (1% and 4% of intermediately methylated epiloci in Dataset 1 and 2, respectively) in normal conditions (Figure 3F, Supplementary Figure 4D), suggesting that intermediate values of average methylation rarely reflect an intermediate methylation amount on different DNA molecules. Instead, intermediate values of average methylation more often reflected the coexistence of fully unmethylated and fully methylated molecules, in presence (D5) or in absence (D3) of intermediately methylated molecules.

The classification of MC profiles to MPs also enabled us to investigate whether epiloci attributed to the different prototype classes were located in genomic regions with different functional characteristics. A shown in Figure 3G and Supplementary Figure 4E, we found that the D1 class was enriched within promoters and exons (chi-square post hoc p-values < 1e-7) and depleted in intergenic regions and introns (chi-square post hoc p-values < 1e-7). On the contrary, the D2 class was mainly located in intergenic regions and introns (chi-square post hoc p-value< 1e-7) and depleted in promoters and exons (chi-square post hoc p-value< 1e-7). Similarly, the D5 class was depleted from promoters and enriched in intergenic regions (chi-square post hoc p-values < 1e-7). We did not find significant differences in the localization of D3 and D4 epiloci.

### Cellular heterogeneity is the strongest contributor to MC profiles

MC profiles recapitulate heterogeneous methylation status among DNA molecules. This heterogeneity can, in principle, reflect both allelic and cellular differences. For most epiloci, these two components cannot be distinguished in a bulk experiment, in which the information on how DNA molecules are paired in individual cells is missing. We reasoned that epiloci present as single copies in the genome could be a good model to investigate the contribution of cellular differences to MC profiles. In fact, at these loci, the presence of multiple MCs can reflect only cellular differences‥

As a first model of DNA regions present as single copies in the genome, we investigated epiloci located on the X chromosome of a male mouse from Dataset 1 (n=1303). For each epilocus, we quantified the cellular heterogeneity in terms of MC counts, i.e. the number of MCs supported by at least 1 DNA molecule. We found MC counts above 1 for most epiloci (>70%), pointing to cellular heterogeneity of DNA methylation as the rule for most epiloci. Of note, the distribution of MC counts’ values for X epiloci did resemble that of autosomal epiloci. This observation seems to suggest that the degree of heterogeneity captured by MC profiles is poorly affected by the copy number of the given epilocus (Figure 4A).

**Figure 4:**
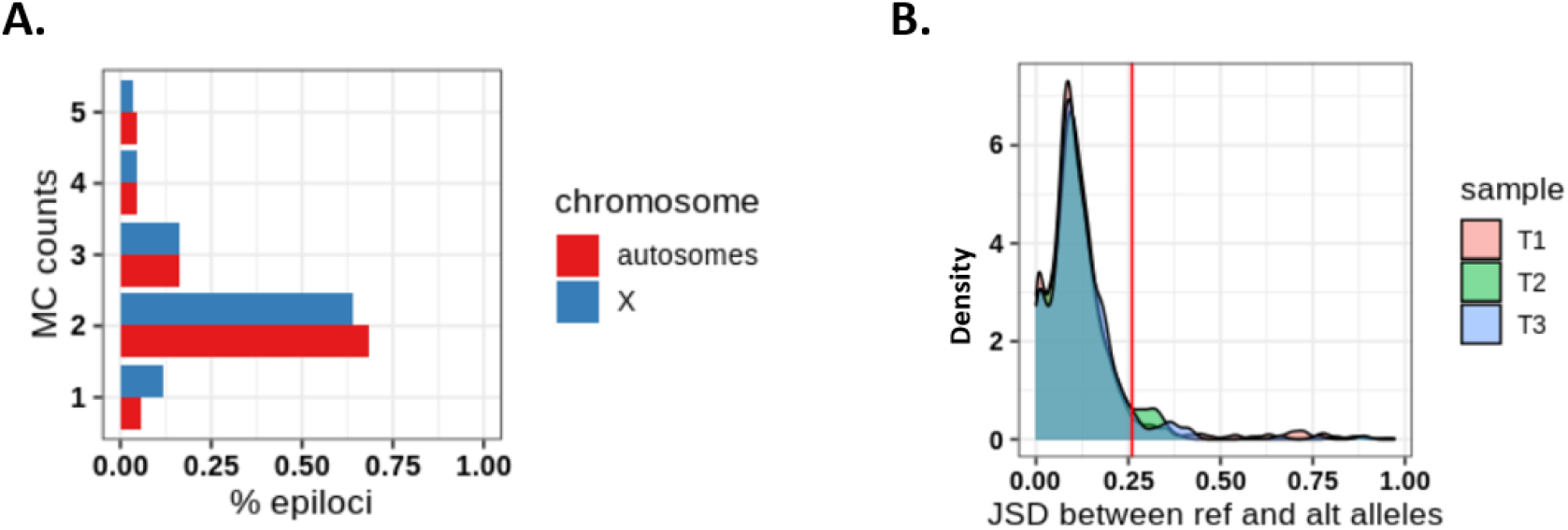
MC profiling of haployd epiloci. **A**: Fraction of epiloci (x-axis) exhibiting a given value of MC count (y-axis) on the X chromosome (blue) and autosomes (red) in a male sample from Dataset 1. **B**: distribution of JSD values between reference and alternative alleles in three samples from Dataset 3.

As an additional model, we studied autosomal epiloci of mice born from two different strains (Dataset 3 in Supplementary Table 1). Based on known polymorphic sites between the two strains, we were able to attribute each read to the respective allele, and to explore the allele specific MC profile for more than 300 autosomal epiloci in 3 mice (see Methods). When analyzing the joint MC profiles, we confirmed the high degree of DNA methylation heterogeneity, with about 95% of the autosomal epiloci having an MC count equal or greater than 1 (Supplementary Figure 6 A-C). However, when directly comparing the MC profiles of the reference and alternative alleles, we found no differences for most of the epiloci, with only a small proportion of epiloci (6-7%) exhibiting allele specific methylation (Figure 4B).

Overall, the results from the analysis of male X chromosome MC profiles and allele specific MC profiles provided evidence for cell-to-cell differences as the major contributor to MC profiles, with evidence of allelic differences only in a small fraction of autosomal epiloci. In addition, we found that the allelic MC profiles of 144 epiloci shared among the samples were mostly stable among sample pairs, thus suggesting that similar patterns of cellular heterogeneity were present in different individuals (data not shown).

### MC profiling individuates a signature of imprinted regions

We tested the capability of MC profiling to discriminate regions undergoing genomic imprinting, a well-known phenomenon of allele specific regulation. In these regions, it is expected that the two alleles differ for their DNA methylation status. Hence, we wondered whether D3 epiloci, in which two pools of molecules exist with opposite DNA methylation status, were enriched at genomic regions flanking imprinted genes.

To test this hypothesis, we assigned each epilocus of Dataset 1 and 2 to its nearest gene (see Methods) and marked the epiloci as associated with imprinted genes if the closest gene was enlisted in Geneimprint (https://www.geneimprint.com/). As shown in Figure 5A and Supplementary Figure 7, the five MPs were differentially represented among epiloci flanking imprinted and not imprinted genes (chi-square test p-values < 2.2e-16). Specifically, epiloci assigned to the D3 pattern were strongly overrepresented among epiloci flanking imprinted genes (chi square post-hoc test p-values < 2e-16), thus confirming that D3 epiloci were preferentially, even though not exclusively, associated with allele specific methylation.

**Figure 5:**
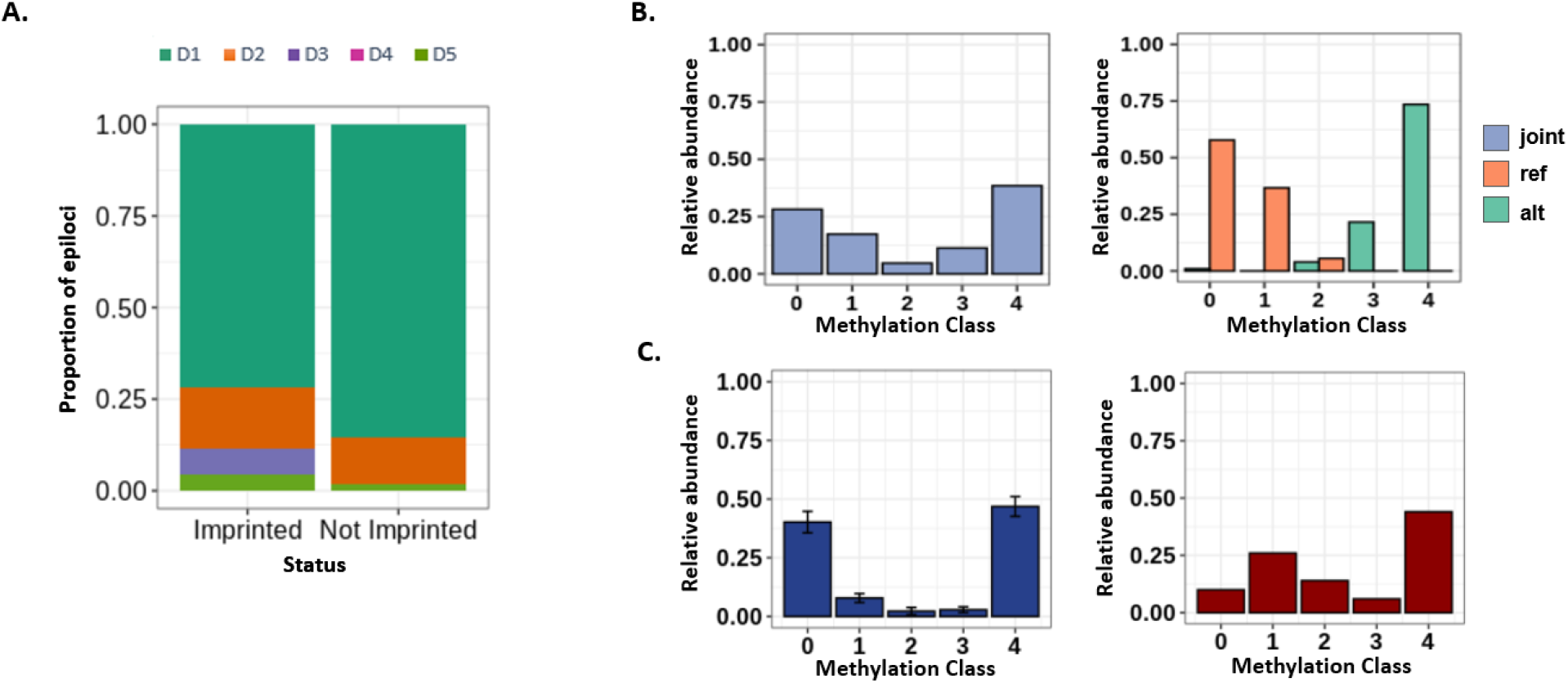
MC profiling of imprinted regions. **A**: proportion of epiloci assigned to the different MPs in imprinted and non imprinted genomic regions. **B**: example of bimodal epilocus flanking the Zdbf2 imprinted gene. The joint MC profile (i.e. the profile obtained without splitting the alleles) is shown in light blue, whereas the profiles of the reference (ref) and the alternative (all) alleles are shown in orange and green respectively. **C**: epilocus in GNAS promoter (chr20:57415288-57415313) with altered MC profile in a tumor sample. For this epilocus, the MC profile averaged on three control samples from Dataset 2 is shown in blue, and the MC profile from the tumor sample from Dataset 4 is shown in red.

As a confirmatory experiment, we searched for D3 epiloci flanking imprinted genes in Dataset 3. We found a single epilocus with these characteristics, located on chr1, upstream of the Zdbf2 imprinted gene. In this locus, differentially methylated regions had been previously described (43). Figure 5B shows how the bimodal joint MC profile at this epilocus results from different profiles on the two alleles, with one skewed towards complete demethylation and the other towards complete methylation. It is worth noting that, for both alleles, MC profiling individuated a certain degree of cellular heterogeneity, since intermediate MCs were also represented.

Loss of genomic imprinting is a well-known epigenetic modification occurring in many tumors (44,45). Considering the association that we found between bimodal D3 MC profiles and imprinted regions, we wondered whether changes in MC profiles of D3 epiloci could be identified in tumor samples. When inspecting epiloci flanking imprinted genes in tumor samples (Dataset 4), we identified an epilocus, located at chr20:57415288-57415313, whose MC profile profoundly changed in one of the samples in respect to controls from Dataset 2. This epilocus was located in the promoter of the GNAS gene, for which loss of imprinting in tumors has been described (Figure 5C) (45).

### MC profiling aids to dissect cell-to-cell differences in DNA methylation on the inactive X

Based on the results obtained from MC profiling of imprinted genes, we decided to investigate whether epiloci located on the X chromosome also exhibited peculiar MC profiles due to the X inactivation process. It is in fact known that, during the inactivation of the X chromosome, most loci are inactivated (subject loci) while others partially or totally escape this inactivation (escapee or variable escapee loci) (46).

We indeed analyzed the MPs for epiloci flanking genes with different inactivation status. First, we assigned X epiloci to the respective MP in two female samples from Dataset 2. Then, we assigned to each epilocus the consensus inactivation status of the nearest gene (46). In this way, we classified 551 epiloci as subject to X chromosome inactivation, 138 as escapee, 56 as variable escapee and 233 as unknown/discordant. As shown in Figure 6A, MPs were represented in different proportions among subject, escape and variable escape epiloci (chi square post-hoc test p-value < 1e-7). Escape epiloci mostly exhibited unmethylated D1 profiles (chi square post-hoc test p-value < 1e-7), whereas subject epiloci mostly exhibited either bimodal D3 or uniform D5 profiles (chi square test post-hoc p-value < 1e-7). Both groups of MPs (D1 and D3/D5) were represented among variable escape epiloci, none of them significantly enriched.

**Figure 6:**
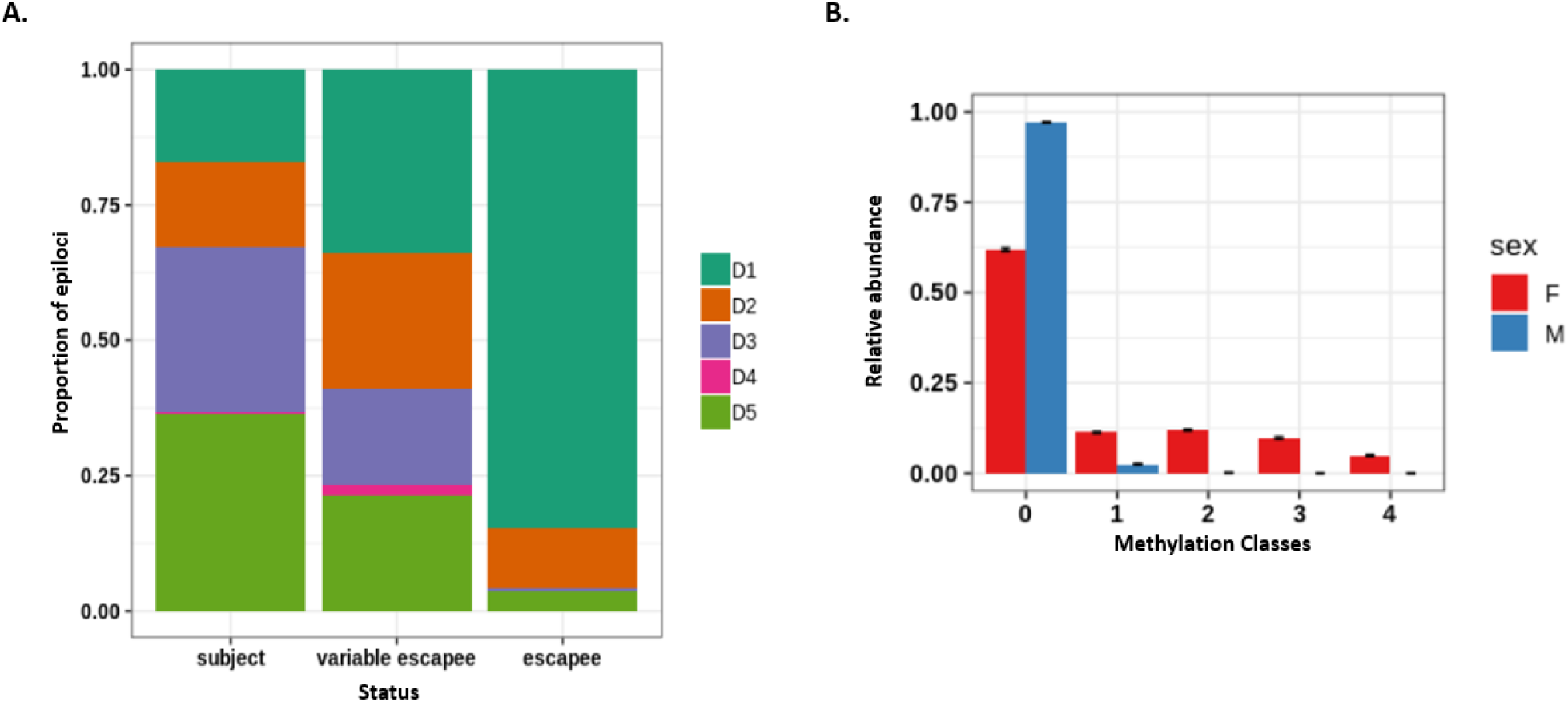
MC profiling of X chromosome epiloci. **A**: classification of epiloci flanking genes undergoing X inactivation (subject), stably escaping X inactivation (escapee), or variably escaping X inactivation (variable escapee); **B**: average MC profile of epiloci classified as D1 in a male sample from Dataset 1(blue) compared to the average profile of the same epiloci in two female samples from the same dataset (red).

We hence decided to investigate the MC profile of the inactive X in Dataset 1, for which two female and one male sample were available. We reasoned that we could deduce the profile of the female inactive X by comparing the MC profile of X epiloci in males and females, and that such deduction would have been particularly feasible for epiloci classified as D1 in the male samples. In fact, in this condition, it could be reasonably inferred that methylated molecules in females mostly resemble the methylation status of the inactive X. We indeed compared the average profiles of 1068 epiloci belonging to the D1 MP in males with the respective average profile in female samples (Figure 6B). The sex difference among the average MC profiles pointed to a heterogeneous DNA methylation status of the inactive X, ranging from being lowly to fully methylated in different cells. Of note, we observed a more gradual methylation status of the inactive X compared to the methylated alleles of imprinted epiloci. This observation is compatible with the previously described discrepancy of average methylation between imprinted and X inactivated genes. In fact, while for imprinted loci one of the alleles is fully methylated, X inactivated genes exhibit partial methylation of the inactive allele (58). In addition, MC profiles suggest that this partial methylation is due to cell-to-cell differences, and not to a partial methylation in all cells.

### MC profiling individuates loci undergoing epigenetic remodeling upon neuronal differentiation

We challenged the ability of MC profiling to capture epigenetic changes among conditions. As a model of epigenetic changes, we choose a dataset of neuronal differentiation.

To this aim, we analyzed MC profiles changes of 115608 epiloci upon differentiation of hippocampal precursors (HP) to granule cells (GC) (Dataset 5). For each epilocus, we calculated the difference of average methylation between differentiated cells and neuronal precursors (delta meth), and quantified the MC profiles’ change by using the Jensen-Shannon distance (JSD). The relationship between these two measures is shown in Figure 7A. The red lines delineate the difference of average methylation observed in 95% of the considered epiloci (0.14), and the black line indicates the JSD threshold (0.26).

**Figure 7:**
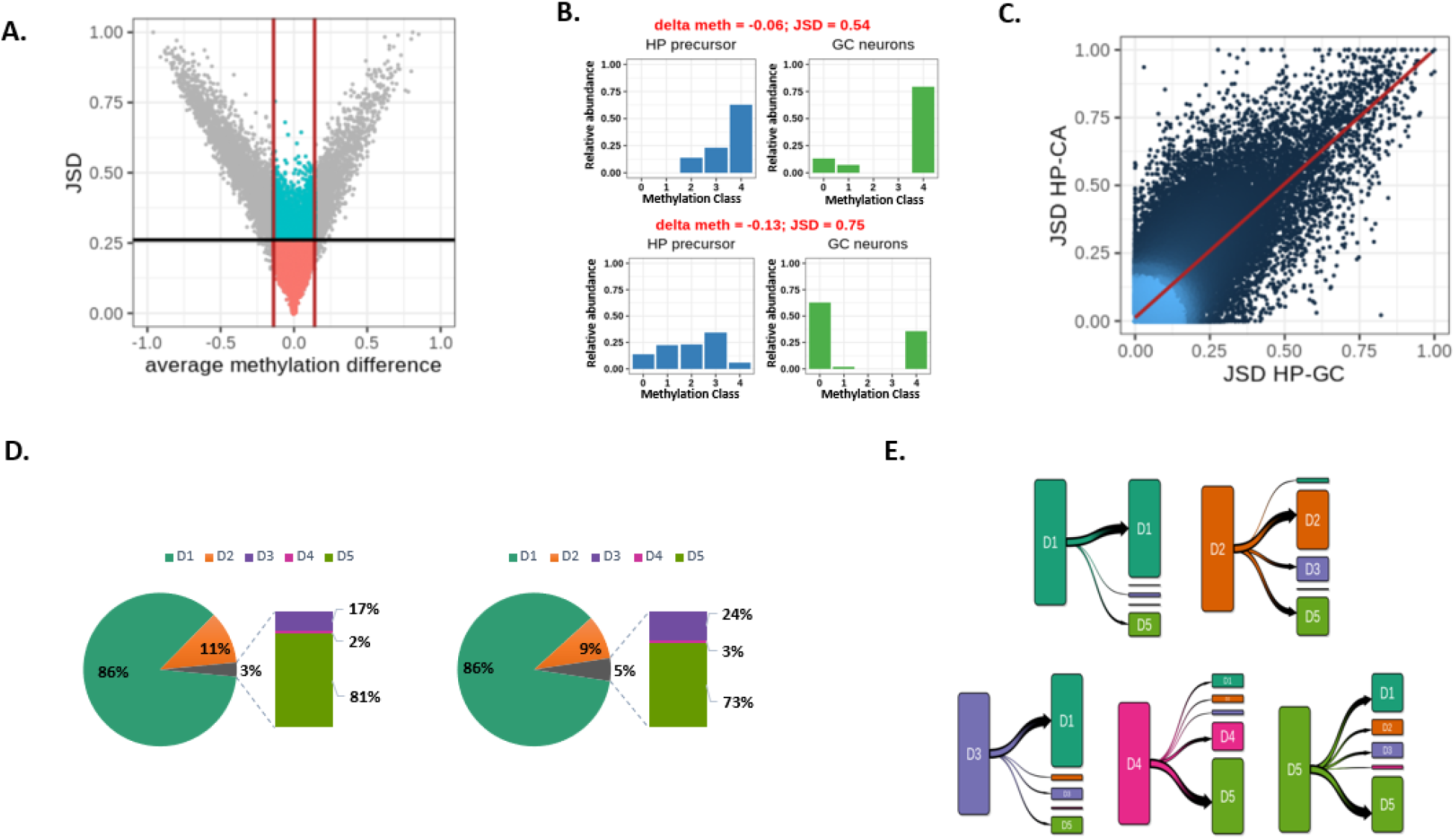
Application of MC profiling on neuronal differentiation. **A**: DNA methylation changes at 104720 epiloci upon differentiation of hippocampal precursors to granule cells. X-axis: average methylation change (delta met); y-axis: MC profile change (JSD). The black line indicates the JSD cutoff, whereas the red lines indicate the 95th percentile of observed delta values. **B**: examples of epiloci with low difference in average methylation but high JSD values between HP and GC MC profiles (epiloci coordinates: chr19:57700749-57700788 and chr1:186924297-186924337). **C**: Comparison of MC profile changes upon differentiation of hippocampal precursors (HP) to granule cells (GC) or CA3 neurons. x-axis: JSD values between MC profiles in HP and GC. y-axis: JSD values between MC profiles in HP and CA. **D**: MPs composition of hippocampal precursors (on the left) and granule cells (on the right) samples. **E**: Transition plot of variant epiloci in the HP-GC pair. For the epiloci assigned to the different MPs in HP cells, the classification in differentiated GC neurons is shown.

As expected, MC profiles’ and average methylation changes were mostly correlated. This relationship strengthened as differences in average methylation approached the maximum, consistent with the fact that huge differences in the amount of methylated cytosines are expected to affect both average methylation and MC profiles. Symmetrically, the relationship between average methylation and MC profiles’ changes weakened for lower values of average DNA methylation and was almost lost below 0.14. In this range, despite a large number of epiloci exhibiting stable MC profiles (n= 104720), a group of 5129 epiloci exhibited significant changes in MC profiles upon differentiation (blue dots in Figure 7A). As examples, Figure 7B shows two epiloci with significant changes of MC profiles and little variation of average DNA methylation. This result suggests that, at these epiloci, MC profiles were remodeled without a significant gain or loss of overall DNA methylation. The results of the enrichment analysis for genes flanking this group of epiloci are shown in Supplementary Figure 8.

To test the association between changes in MC profiles and the process of neuronal differentiation, we checked the consistency of the MC profiles changes upon differentiation of the same precursor in a different type of neuron. Notably, we found a high correlation between MC profiles changes for the 97119 epiloci examined upon differentiation of hippocampal precursors to granule cells or CA neurons (Pearson R 0.81, Figure 7C), according to the previously described high similarity among these differentiation processes (47).

To further establish the relationship between the changes in MC profiles and epigenetic remodeling upon cell differentiation, we explored the chromatin landscape, summarized by chromHMM labels, associated with the analyzed epiloci. We observed that MC profile changes more probably involved epiloci located in regions that also underwent chromatin changes upon differentiation (Fisher test p-value < 2.2 e-16). In fact, only 5% of epiloci located in genomic regions with stable chromatin marks underwent changes in their MC profile, whereas 22% of epiloci undergoing chromatin changes also changed their MC profile, thus suggesting that our approach was probably identifying loci undergoing epigenetic remodeling.

When investigating the genomic localization of developmentally variant epiloci discovered by our approach, we found that they were slightly depleted outside CpG islands and promoters (Fisher test p-values < 2.2 e-16) both in HP-GC and HP-CA transitions.

Being JSD is a symmetric distance, it only quantifies the dissimilarity between two MC profiles, but does not return the information on whether this dissimilarity corresponds to a gain or loss of DNA methylation. Thus, we turned to the analysis of prototype classes to qualitatively interpret MC profile changes upon differentiation. The prototype class composition for HP and GC is shown in Figure 7D.

First, we asked whether changes were occurring at epiloci exhibiting peculiar MC profiles in neural precursors. We found that epiloci classified as D1 remained mostly stable, whereas epiloci assigned to the other classes mostly changed their MC profile upon differentiation (chi-square post-hoc p-values < 1e-7).

We then analyzed the prototype class composition in differentiated neurons, and found a depletion of D2 epiloci and an increased fraction of D5 epiloci (chi-square post-hoc p-values < 1e-7), suggesting that the methylated status in differentiated neurons tends to be more heterogeneous among different cells.

Finally, to better characterize how MC profile changes were occurring, we analyzed the prototype class transitions upon differentiation. In Figure 7E, for each prototype class in neuronal precursors, we show the final prototype class in differentiated neurons.

We noticed that for a consistent fraction of D1 and D2 epiloci, MC profiles’ changes did not correspond to class transitions, meaning that these epiloci were shifting toward a higher or lower DNA methylation heterogeneity. We also noticed that a reduced fraction of epiloci evolved toward the D2 class upon differentiation.

Interestingly, most D3 epiloci evolved to lower methylation upon differentiation, transiting to the D1 class. Thanks to prototype class analysis, we could interpret this demethylation as a negative selection of the fully methylated molecules that were present in neural precursors.

All together, these results indicate that our approach well captures quantitative and qualitative DNA methylation changes upon neuronal differentiation that might be underestimated or overlooked by an average methylation based approach.

## DISCUSSION

Each cell is a uniqum. Evidence has accumulated that even in morphologically homogeneous cell populations extensive differences can be highlighted at multiple molecular levels, and that these differences are relevant to biological processes (31).

Single-cell DNA methylation assays promise to be the standard technique to study DNA methylation heterogeneity in cell populations (30,48). However, single-cell DNA methylation technologies still generate very sparse data, in a limited number of cells per sample, and at high cost (28,32). Alternatively, cell-to-cell differences can be deduced by studying the methylation patterns of consecutive cytosines in sequenced reads from bulk experiments, assuming each read coming from a single DNA molecule (19,28,33–36). This approach can stand comparison with single-cell assays when the goal is to obtain a robust description of DNA methylation of a genomic region in a cell population (28).

Building on top of our experience on deep targeted bisulfite sequencing, in this study we propose MC profiling as a genome-wide approach to the study of DNA methylation heterogeneity. Given an epilocus holding 4 CpG sites, we defined its MC profile as the ensemble of the relative abundances of molecules sharing an equal number of methylated cytosines (Methylation Classes, MCs). Such an approach, while incorporating information on the overall average methylation of a region, directly informs on the different methylation levels, and their abundance, observed in a pool of molecules. This information is usually not, or poorly, taken into account by other approaches, which directly quantify the degree of cellular heterogeneity through the analysis of individual arrangements of methylated cytosines in single DNA molecules (epialleles).

A previous study showed that the methylation level of individual molecules can be used to adjust mean methylation indices for cell heterogeneity, thus improving prediction of gene expression levels compared to the overall average methylation (35). This method quantifies DNA methylation at promoter regions as the ratio of reads holding ≥1 methylated cytosines to the total number of reads mapped to the promoter. MC profiling is in line with this logic, but further enlarges the information on DNA methylation heterogeneity by considering molecules with different methylation levels as separate entities.

A conceptually similar approach to MC profiling has been proposed in (38–40). In these studies, DNA methylation is expressed as the probability mass function (PMF) of methylation levels that could be observed in a pool of molecules, which resemble the concept of our MCs. This approach, specifically designed to deal with the low coverage of WGBS experiments, has provided novel insights on DNA methylation heterogeneity and its disposition across the genome, its evolution upon differentiation, aging and cancer, and its relationship with the genetic background (38–40). The biggest difference between this approach and MC profiling is that while the PMF is predicted from a mathematical model applied to DNA methylation data, the frequencies of MCs are empirically estimated from experimental data.

To quantify the dissimilarity between MC profiles, we adopted the Jensen-Shannon distance (49). This dissimilarity measure has been applied in bioinformatics and epigenetics (38–40,50–52).

To set the parameters of our approach, we synthesized low coverage 4 CpG datasets from an in-house database of high-coverage amplicon bisulfite sequencing data (27,53–55). In this context, we provided a systematic quantification of the impact of coverage on the accuracy of MC profiles, and estimated the expected error associated with MC profiles at a coverage of 50 reads. In our opinion, these results could serve as guidelines to orient qualitative analysis of DNA methylation in low coverage settings.

Here, similarly to previous studies (41,42), we adopted a classification procedure, assigning each MC profile to the most similar among 5 reference profiles. This classification scheme provided us an interpretable representation of each MC profile. Furthermore, it provided us with a qualitative property to be compared across epiloci. Finally, being this a fixed scheme, we could apply it and directly compare the results on different conditions and species.

We demonstrated that MC profiles were stable among different samples and neighboring epiloci. Previous studies illustrated that DNA methylomes exhibit high inter-individual stability, especially in CG dense regions (56,57). Concordant epigenetic marks, including DNA methylation, across genomic blocks have been also described (23,38,58). Altogether, our results are in line with previously described patterns of regional and inter-individual stability of DNA methylation, and suggest that MC profiles capture controlled DNA methylation dynamics rather than stochastic fluctuations of methylation levels.

In this paper, we applied MC profiling to gain insights on methylation heterogeneity in various biological contexts.

Firstly, we profiled regions either present in single copies in the genome of individual cells or carrying heterozygous polymorphisms that enabled distinguishing the two alleles. We found that cell-to-cell differences were the strongest contributor to the molecular heterogeneity incorporated in MC profiles. Allelic differences contributed only in about 6% of analyzed regions, in agreement with the fraction of allele specific methylation described in most studies (39,59,60).

Secondly, we tested the capability of MC profiling to inspect known examples of mono-allelic regulation, i.e. genomic imprinting and X-inactivation, in which DNA methylation is notably involved.

When we analyzed the MC profiles of epiloci located in proximity of known genomic imprinted regions, we found that bimodal MC profiles were overrepresented. This was expected, considering the known opposite methylation pattern of the two parental alleles at imprinted regions (61). For an epilocus located upstream of the Zdbf2 gene, holding a polymorphic site, we were able to clearly show opposite MC profiles on the two alleles.

Loss of imprinting (LoI) has been described in several tumors (44,45). As a proof of concept, we showed that MC profiling can capture LoI in a leukemic sample, suggesting that this approach could be adopted in this field. We illustrated an example of MC profile alteration in the promoter of GNAS, which LoI has been described in diverse types of cancer (45). Consistent with a previous study, we found that the MC profile was altered toward gain of DNA methylation in the tumor sample (45). In addition, MC profiling suggested that this gain was not homogeneously accomplished in the whole cell population.

We then analyzed the MC profiles of epiloci located on the X chromosome in female samples. First, we compared MC profiles of epiloci flanking genes with reported differential inactivation status. Consistent with previous findings (62–64), escapee epiloci showed homogeneous DNA methylation on both X copies, being unimodally fully methylated or unmethylated. Subject epiloci, on the contrary, were enriched for more heterogeneous MC profiles (D3 and D5), compatible with different DNA methylation status of the two alleles (62–64).

To further inspect the DNA methylation status of the inactive X, we selected the X epiloci with a fully unmethylated profile in male samples, and examined the corresponding MC profiles in female samples to infer the profile of the inactive X. We showed a prevalence of intermediately methylated classes on the inactive X, accompanied by high cellular heterogeneity. Incomplete DNA methylation of the inactive X was described in (65) at single CpG level, thus marking a difference between the X inactivation and the genomic imprinting processes that was well reflected in our analysis. It is worth noting that the prevalence of intermediately methylated MCs that we found with our approach also suggested a difference between the methylated status on the inactive X and at autosomal epiloci, suggestive of peculiar mechanisms intervening in DNA methylation establishment and regulation on the inactive X.

The methylation status of the inactive X appeared also to be highly heterogeneous among different cells. We speculate that this cellular epipolymorphysm could almost in part find its reflection in differences of X inactivation status between equivalent cells described in single-cell RNA-seq studies (66,67).

Finally, we applied MC profiling to the analysis of DNA methylation changes in different conditions. In particular, we examined profiles’ changes upon differentiation, when epigenetic remodeling is expected to occur. We adopted the Jensen-Shannon distance to capture epiloci with significant differences in MC profiles between neural precursors and differentiated neurons. Since JSD is a symmetric distance measure, it didn’t return the information on whether MC profiles changes correspond to gain or loss of DNA methylation. Thus, we examined the pattern transitions to gain insights on how profiles’ changes were occurring. Combining the analysis of JSD and pattern transitions provided us a comprehensive picture of DNA methylation differences among conditions: in fact, we could distinguish profiles changes associated with unvaried patterns (and thus, with stable reciprocal proportion of DNA molecules with different methylation levels) from profiles changes accompanied with pattern transitions (which indicate a redistribution of the proportions of molecules with different methylation levels).

As expected, we found that MC profile changes captured by JSD correlated with average DNA methylation gain or loss at most epiloci. However, we described MC profile changes at almost constant DNA methylation for more than 5000 epiloci. Qualitative DNA methylation changes occurring with little to no changes in overall average methylation were also described in (38–40), indicating that such an approach can be even more informative than the average methylation-based approach in the analysis of dynamic systems.

Interestingly, we found that MC profile changes were enriched at CpG islands, which were described to be spared from most epigenetic changes in the original study (47). The association that we found with changes of chromatin marks, as well as the concordance of MC profiles changes upon differentiation in two different neuronal subtypes, points to exclude random variations occurring at these epiloci. Instead, considering that most epiloci exhibited stable prototype classes in precursors and differentiated neurons, it is possible that MC profiling has captured changes in cellular heterogeneity that were overlooked by the average methylation-based approach.

Applying MC profiling to RRBS data can give insights on cellular epigenetic heterogeneity from plenty of already available datasets in public repositories. However, it strongly limits the analysis to CpG islands and immediately proximate regions (15,16), which are usually reported to be resistant to DNA methylation and to exhibit low regulatory plasticity in normal conditions (3,68,69). This limit is further exacerbated when selecting target regions harboring 4 CpGs (the epiloci of this study). The required coverage of 50 reads strongly limits the applicability of the proposed approach to Whole Genome Bisulfite Sequencing (WGBS) data. However, more unbiased enrichment assays have been developed which combine high throughput sequencing with selection of target regions through PCR or capture-based trapping (70,71). that are natively less biased toward CG dense regions and could fit our coverage requirements of MC profiling. We indeed believe that applying MC profiling to these experiments could further extend our observations outside CG dense regions.

## CONCLUSIONS

In this study, we have presented a novel approach named MC profiling aimed at exploring DNA methylation heterogeneity at multiple target regions in bulk bisulfite sequencing experiments. MC profiling is built on the concept of MCs, groups of molecules holding the same number of methylated cytosines. DNA methylation is indeed represented through MC profiles, i.e. the relative abundances of possible MCs for a given region.

MC profiles directly incorporate the average methylation of a given region, and inform on how it is contributed by single DNA molecules. Thus MC profiles offer a functional view of DNA methylation heterogeneity in a sample. MC profiles are empirically estimated from sequencing reads, and are independent on a priori parametrization of DNA methylation dynamics. Moreover, MC profiles retain all information from a pool of molecules, and enable the direct visualization of DNA methylation heterogeneity of a given region.

In this study, we showed that MC profiling identified cell-to-cell differences as the prevalent contributor to DNA methylation heterogeneity, with allele differences emerging in a small fraction of analyzed regions. Moreover, MC profiling led to the identification of signatures of loci undergoing genomic imprinting and X inactivation, and highlighted differences between the two processes. When applied to a dynamic system, MC profiling identified DNA methylation changes in regions with almost constant average methylation. Altogether, our results indicate that MC profiling can provide useful insights on the epigenetic status and its evolution at multiple genomic regions.

## METHODS

### MC profiling

#### a. Epilocus definition and MC profiling

We analyzed the methylation status of regions holding 4 CpG sites, with the first and the fourth CpGs delimiting the region. We refer to these regions as epiloci in the text. For a certain epilocus, we counted the different arrangements of methylated and unmethylated cytosines found in sequencing reads. We then grouped the observed arrangements in 5 methylation classes (MCs) according to the number of methylated cytosines they bear. Hence, we computed the epilocus MC profile as the fraction of reads supporting a given MC out of the total number of reads. Only reads spanning the entire epilocus were considered in the computation of the corresponding MC profile.

#### b. Measure of dissimilarity between MC profiles

We adopted the Jensen-Shannon Distance as a measure of dissimilarity between two MC profiles. The Jensen Shannon Distance (JSD) quantifies the degree of dissimilarity between discrete distributions P_1_ and P_2_ (49), and is defined as

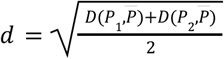

where 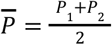 represents the average distribution of two MC profiles, and
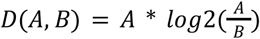 is the Kullback-Leibler divergence between the two MC profiles (72).

#### c. Establishment of MC profiling thresholds

We used deep amplicon bisulfite sequencing (D-ABS) data to synthesize 4-CpG low coverage datasets, starting from an in-house database of D-ABS amplicons produced in previously published studies (27,53–55). Detailed descriptions of the employed amplicons can be found in Supplementary Table 2.

First, we split each amplicon into non-overlapping regions made up of 4 CpGs, thus obtaining several 4-CpG high-coverage datasets. Among these datasets, we selected those with higher coverage (number of reads > 20000). Since we expect that fully methylated or unmethylated profiles would be better captured at low coverage than intermediately methylated ones, due to the higher number of methylation classes with non-zero abundance, we selected 4-CpG datasets with average methylation levels spanning the entire range from 0 to 1 and enriched for datasets with intermediate average methylation. In this way, we selected 25 datasets, representative of 5 groups according to the average methylation level (Supplementary Table 3-4).

To simulate low coverage datasets, we randomly sampled a fixed number of reads from each 4-CpG dataset.

We adopted the low coverage datasets to address the following issues:

1) the minimum coverage to minimize the error between a reference MC profile (i.e., the MC profile computed from the high coverage dataset) and an estimated MC profile (i.e., the MC profile computed from a low coverage dataset)

To address this point, we synthesized 1000 low coverage 4 CpG datasets for coverage values ranging from 20 to 200. From each dataset, we calculated the MC profile, and computed the JSD from the MC profile of the respective 4 CpG high-coverage dataset. As shown in Supplementary Figure 1A, the JSD values decreased as the coverage increased, as expected. In particular, JSD values dropped between 25 and 50 reads (Supplementary Figure 9A). A similar gain in accuracy is achieved by triplicating the coverage (i.e. achieving a read number higher than 150). Based on these observations, we considered a region covered by at least 50 reads to be eligible for MC profiling.

2) the minimum value of JSD to consider 2 MC profiles as different.

To address this point, we simulated 1000 pairs of low-coverage datasets with a fixed coverage of 50 reads. For each dataset pair, we computed the JSD among the estimated MC profiles. Ideally, two read groups sampled from the same dataset should exhibit very similar, if not identical, profiles, with a JSD value approaching 0. In practice, however, the estimated profiles differed to a certain extent. As shown in Supplementary Figure 9B, at a coverage of 50 reads, MC profiles exhibited a JSD lower than 0.26 (min=0.22, max=0.28) for 95% of the experiments. We concluded that MC profiles having a JSD higher than 0.26 could be defined as different with an error equal or lower than 0.05. Hence, when comparing two MC profiles, we considered them to be different when we observed a JSD above 0.26.

#### d. Epilocus filtering and multiple samples handling

In individual samples, epiloci with coverage lower than 50 reads and higher than 99th percentile were filtered out. Overlapping epiloci were also removed.

To handle MC profiles observed in different samples at a given epilocus, we first computed the JSD between the possible sample pairs and retained those epiloci with JSD below 0.26 in all the pairs. For these epiloci, we computed the average MC profile by averaging the relative abundance of each MC among the samples.

In this way, we obtained a consensus MC profile representative of all samples in a given condition. This enabled us to directly compare consensus profiles of an epilocus in two conditions through the JSD.

#### e. MC profiles classification

To provide biological interpretation of MC profiles, we adopted a data compression scheme, assigning MC profiles to 5 methylation patterns (MP). We first defined 5 reference profiles, from D1 to D5, here referred to as prototypes (Figure 2D). To assign an MC profile to the nearest MP, we computed its JSD from the 5 prototypes, and assigned it to the MP corresponding to the prototype with minimum JSD. To check the appropriateness of our classification procedure, we compared the JSD of each MC profiles from the two nearest MPs, with the lower JSD value representing the distance from the membership pattern centroid (Within Class Distance, WCD) and the second value representing the distance from the nearest outer pattern centroid (External Class Distance, ECD). We observed that the ECD-WCD ratio exceeded 1.5 for 98% of MC profiles in Dataset 1 and 95% of MC profiles in Dataset 2 (Supplementary Figure 10). Thus, we concluded that the proposed scheme was roughly consistent and representative of the diverse MC profiles observed in our datasets.

### Datasets

We analyzed previously published RRBS data and enhanced RRBS data (47,59,73,74). The data were publicly available in the GEO database (https://www.ncbi.nlm.nih.gov/geo/) with the following accessions: GSE66121, GSE130735, GSE53714, GSE72700. When a consistent discrepancy existed for the number of epiloci covered by at least 50 reads in different samples, we retained those samples providing the highest number of epiloci. A detailed description of the samples adopted from each dataset is included in Supplementary Table 4.

### Data processing

#### a. RRBS raw data processing

Raw RRBS data were processed using an in-house pipeline. Fastq files were first quality checked by using FastQC (https://www.bioinformatics.babraham.ac.uk/projects/fastqc/). Low-quality bases were removed using Trim Galore v0.6.6 with parameters --rrbs and -paired for paired end experiments (https://www.bioinformatics.babraham.ac.uk/projects/trim_galore/). The obtained reads were aligned to the hg19 or mm10 reference genomes by using Bismark v0.23.0 with default parameters (75). The obtained BAM files were then sorted and indexed using the SAMtoolsKit (http://www.htslib.org/).

#### b. Deep – Amplicon Bisulfite Sequencing data processing

D-ABS data were processed as previously described (27,53,54). In brief, paired-end reads were merged in a single fastq file by using PEAR (minimum overlapping residues equal to 40) (https://cme.h-its.org/exelixis/web/software/pear/doc.html). The fastq file was then converted to fasta by using PRINSEQ (http://prinseq.sourceforge.net/).

#### c. Epiallele counts extraction

For RRBS data, epiallele counts were extracted from BAM files using the *EpiStatProfiler* R package (https://github.com/BioinfoUninaScala/epistats). First, genomic regions covered by at least 50 reads were individuated through the *filterByCoverage* function. Target regions holding 4 CpGs (the epiloci described in this manuscript), stepping up to 1 CpG at time, were then defined by using the *makeBins* function. The maximum length of the target regions was set from 70 to 100 bps, depending on the specific library design. Epiloci covered by at least 50 reads spanning the entire region were retained. Finally, selected epiloci were analyzed by using the *epiStatAnalysis* function with default parameters. For each epilocus, the function returns a table with summary statistics (including the average methylation), and a file with epiallele counts. This latter was then analyzed through in-house R scripts to compute the epilocus MC profile, as described above.

For D-ABS data, epiallele counts were then extracted by using the AmpliMethProfiler tool (76). The MC profile of the amplicon was then computed following the same procedure of RRBS epiloci.

#### d. Allele-specific alignment sorting

To perform allele specific MC profiling, we applied the pipeline based on the SNPsplit tool (77) on a dataset of crossed strain mice. First, the positions holding alternative sequences between the strains were extracted from the VCF file downloaded from the Mouse Genomes Project repository (ftp://ftp-mouse.sanger.ac.uk/current_snps/mgp.v5.merged.snps_all.dbSNP142.vcf.gz), and were masked from the reference mm10 genome by using the SNPsplit_genome_preparation function in single strain mode. Fastq files were then aligned to the masked genome by using Bismark 0.23.0 with default parameters. The reads aligned to polymorphic sites were assigned to the respective allele by using the SNPsplit function. In brief, the reads aligned to variant positions were tagged (SNPsplit-tag internal function), assigned to the reference or to the alternative allele (tag2sort internal function), and written down in separate bam files. We ran the SNPsplit function in --bisulfite mode to automatically discard the reads aligned to C/T or T/C variants on the forward strand and to G/A or A/G variants on the reverse strand, since these variants cannot be distinguished from a methylation status call. The bam files relative to the reference and the alternative allele were processed independently with the EpiStatProfiler tool to obtain the epiallele counts and to compute the MC profile. At the end, we were able to profile 2749, 460, 314 autosomal epiloci in three mice, with a minimum coverage of 50 reads on both alleles.

#### e. Epiloci annotation

Epiloci were annotated by using the *annotatr R package* against hg19 and mm10 CpG tracks and hg19 and mm10 genes tracks.

Epiloci were associated with the nearest genes by using the *seq2pathway* R package. To minimize the number of genes associated with an epilocus, the ‘adjacent’ parameter was adopted, thus enabling to assign each epilocus to the closest genes only. For the association, the *FullResult* output was considered, which also included non-coding genes.

To test the association between MC profiles and chromatin marks changes, epiloci were annotated using the chromHMM segmentation tracks for mouse hindbrain (E10 and P0) downloaded from UCSC (78). Each epilocus was annotated with the label of the genomic segment with the highest overlap. The E10 hindbrain track was used to annotate epiloci of hippocampal precursors, whereas the P0 hindbrain track was adopted to annotate epiloci in Granule cells and CA neurons.

### Statistical analysis

#### a. Classification concordance of neighboring epiloci

To test the classification concordance of neighboring epiloci, we first binned the genome into 1 kb long regions. We then intersected the bins’ coordinates with that of epiloci shared by all the samples in the dataset (see Epilocus filtering and multiple samples handling). We removed the bins harboring less than 3 epiloci and labeled the remaining ones as concordant if they hold epiloci assigned to the same prototype class, and discordant otherwise. We tested the hypothesis that the number of concordant bins was higher than the one expected by chance by bootstrapping. In brief, we scrambled the epiloci grouped in each bin, such that the overall number of bins together with the number of epiloci they hold reflected those observed in experimental data, but the epiloci were no longer grouped in a bin based on their proximity but were randomly sampled without replacement from the dataset. We repeated this procedure 1000 times, and each time we counted the number of scrambled bins classified as concordant. We thus obtained the distribution of the number of concordant bins expected by chance, which we compared with the number of concordant bins observed in experimental data.

#### b. MC profiles heterogeneity

For haploid models, we adopted the epilocus MC counts, i.e. the number of MCs with non-zero relative abundance, to estimate the degree of cellular heterogeneity of DNA methylation. The MC counts distribution of haploid epiloci was compared to that of dyployd epiloci to estimate the contribution of allelic heterogeneity to MC profiles. For male X epiloci, autosomal epiloci in the same sample were used as dyployd reference, whereas for polymorphic epiloci the joined MC profiles were used. When directly comparing the MC profiles of an epilocus on the two alleles in the second model, we controlled for coverage differences. We found no significant differences between the two alleles (paired Wilcoxon test p-values < 0.01).

#### c. Statistical test

All the statistical analyses were performed using R software (version 4.0) with an alpha value set for p < 0.01.

Association between categorical variables was tested for statistical significance trough Fisher test (when both categorical variables had two possible values) and through chi-square test and post-hoc analysis of chi-square residuals (*chi.square.posthoc.test* function from the homonymous R package, adopting Bonferroni correction to control for alpha inflation).

Difference between mean distance among epiloci in concordant and discordant bins was tested through the Mann-Whitney test.

Enrichment analysis for 5129 epiloci with significant changes in MC profiles and stable average methylation upon differentiation was performed using GREAT version 4.0.4 (79), using default parameters and the coordinates of all the analyzed epiloci (115608) as background.

## DECLARATIONS

### Availability of data and materials

The datasets analyzed in this study are publicly available in GEO with accessions GSE66121, GSE130735, GSE53714, GSE72700.

The R code utilized is available on GitHub (https://github.com/EpigenomicsLAB/MCprofiling_code.git).

