## Supplementary Material for "MC profiling: a novel approach to analyze DNA methylation heterogeneity from bulk bisulfite sequencing data"

### *Supplementary Figures*

**Supplementary Figure 1: Number and genomic distribution of analyzed epiloci. A:** number of epiloci in three samples from Dataset 1 (M1:3) and Dataset 2 (H1:3). B: annotation of epiloci in respect of CpG density and genic regions in three samples from Dataset 1 (M1:3) and Dataset 2 (H1:3).

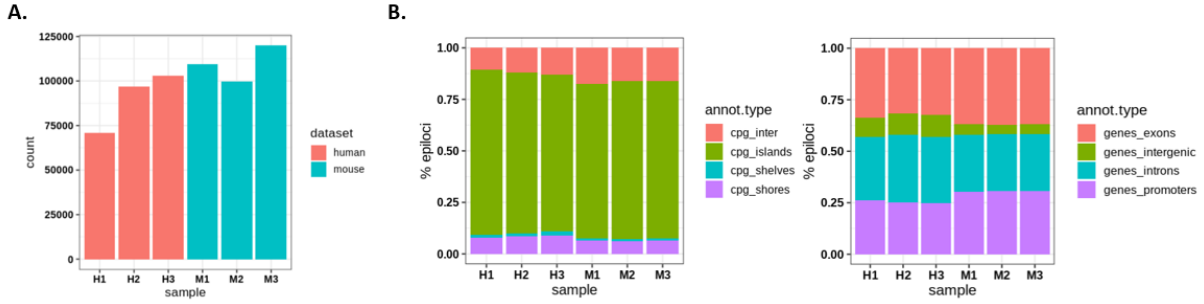

**Supplementary Figure 2: Fraction of genomic functional regions holding at least 1 epilocus in Dataset 1 (panel A) and Dataset 2 (panel B) samples**

**A.**

| | CGI (17017) | Shores (32080) | Shelves (29037) | Promoters (142446) | Exons (841916) | Introns (699470) | Intergenic (29293) |
|---|---|---|---|---|---|---|---|
| M1 | 75% | 13% | 3% | 25% | 6% | 7% | 15% |
| M2 | 74% | 12% | 2% | 25% | 6% | 6% | 13% |
| M3 | 77% | 14% | 3% | 27% | 7% | 7% | 15% |

**B.**

| | CGI (28691) | Shores (51914) | Shelves (43747) | Promoters (82960) | Exons (742493) | Introns (659327) | Intergenic (17027) |
|---|---|---|---|---|---|---|---|
| H1 | 49% | 8% | 2% | 23% | 4% | 5% | 17% |
| H2 | 58% | 11% | 3% | 28% | 5% | 6% | 22% |
| H3 | 58% | 13% | 3% | 28% | 5% | 6% | 23% |

**Supplementary Figure 3: Average DNA methylation of epiloci assigned to the different MPs in Dataset 1 (first row) and Dataset 2 (second row) samples**

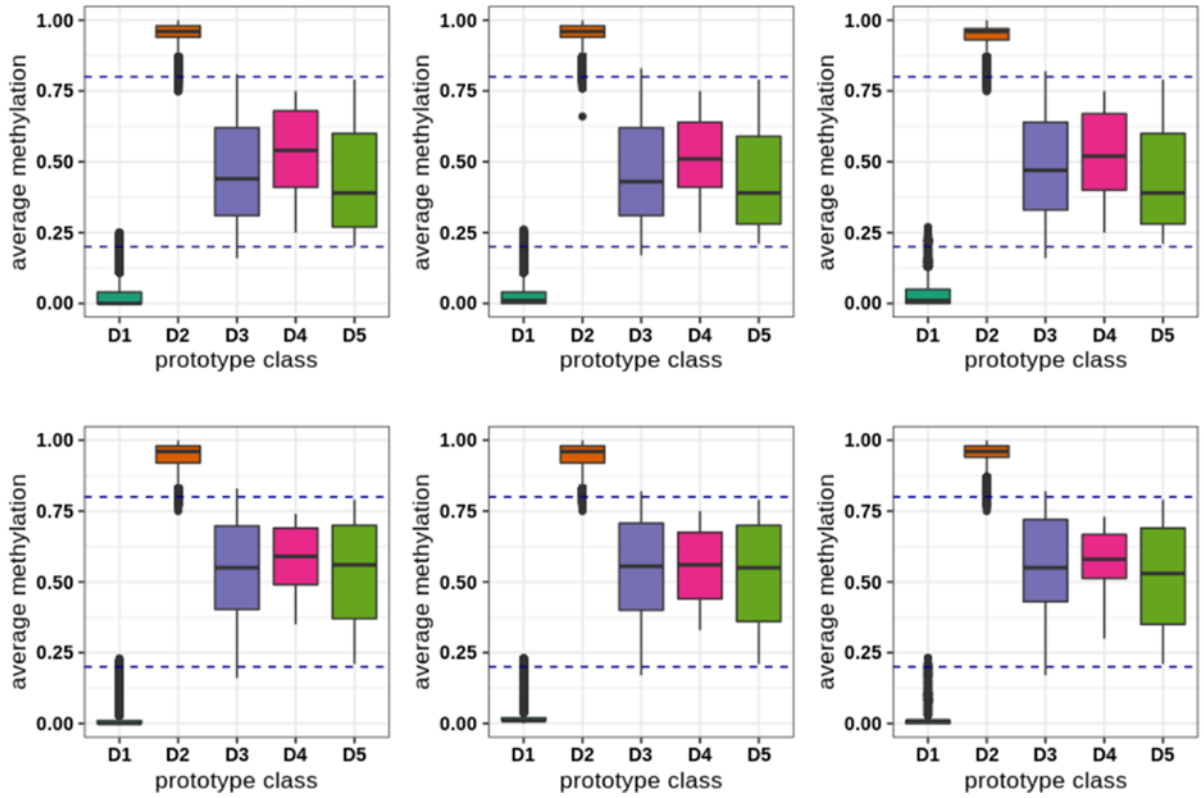

***Supplementary Figure 4: MC profiling results for Dataset 2.*** **A:** Distribution of MC profile distance between sample pairs. The red line indicates the cutoff of JSD. **B:** Genomic annotation of epiloci with stable or variant MC profiles. **C:** average distance between epiloci inside concordant and discordant bins (Wilcoxon test p-value 5.624e-13). **D:** Fraction of epiloci attributed to the different MPs. **E:** genomic annotation of epiloci assigned to the different MPs.

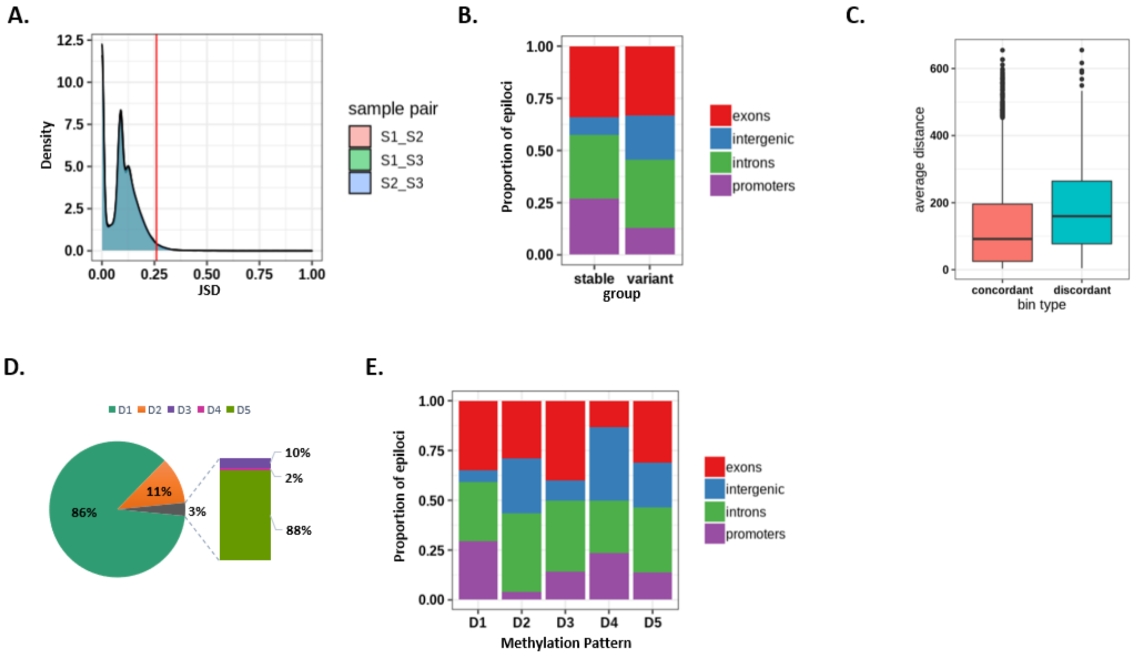

**Supplementary Figure 5: Distribution of concordant bin numbers resulting from bootstrapping analysis in Dataset 1(A) and Dataset 2(B).** In both panels, the red line indicates the observed value

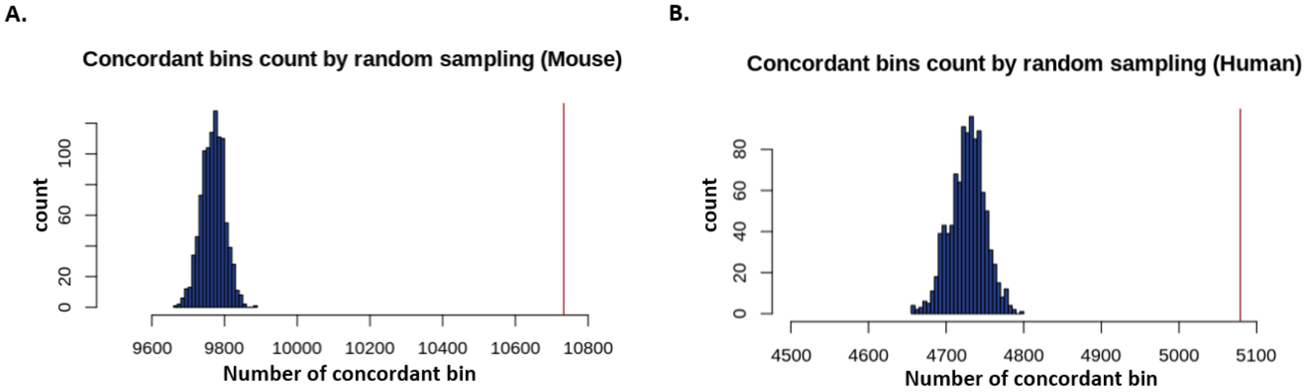

**Supplementary Figure 6: Contribution of cellular heterogeneity to MC profiles at polymorphic epiloci.** Molecular heterogeneity of joined MC profiles in three samples from Dataset 3. The y-axis indicates the MC count value, i.e. the number of MCs with non-zero relative abundance. The x-axis indicates the proportion of epiloci with a given MC count.

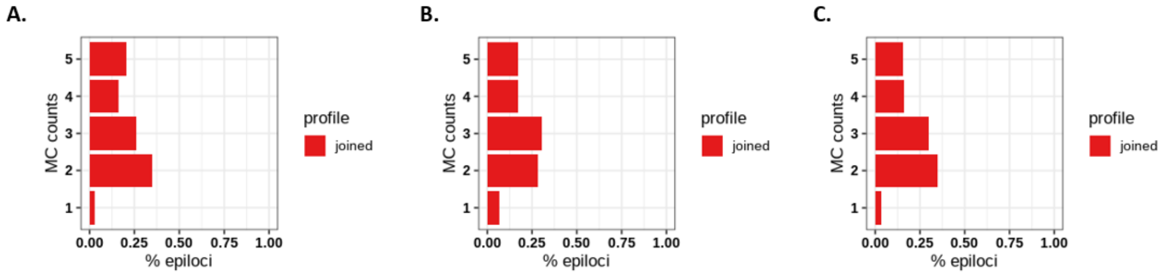

**Supplementary Figure 7: Proportion of epiloci assigned to the different MPs in imprinted and non imprinted genomic regions in the Dataset 2.**

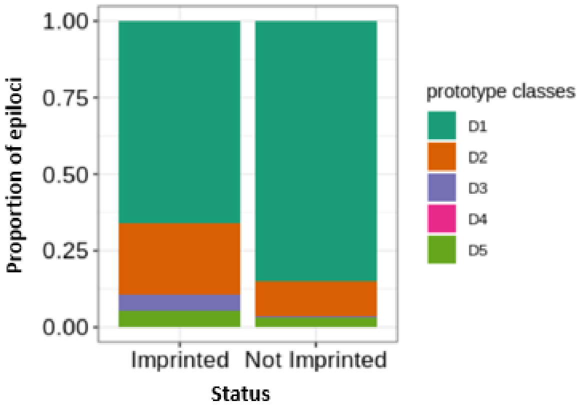

**Supplementary Figure 8: Gene set enrichment analysis for genes associated with 5129 epiloci with changes in MC profiles and stable average methylation upon differentiation.** Genes associated with all analyzed epiloci were used as background.

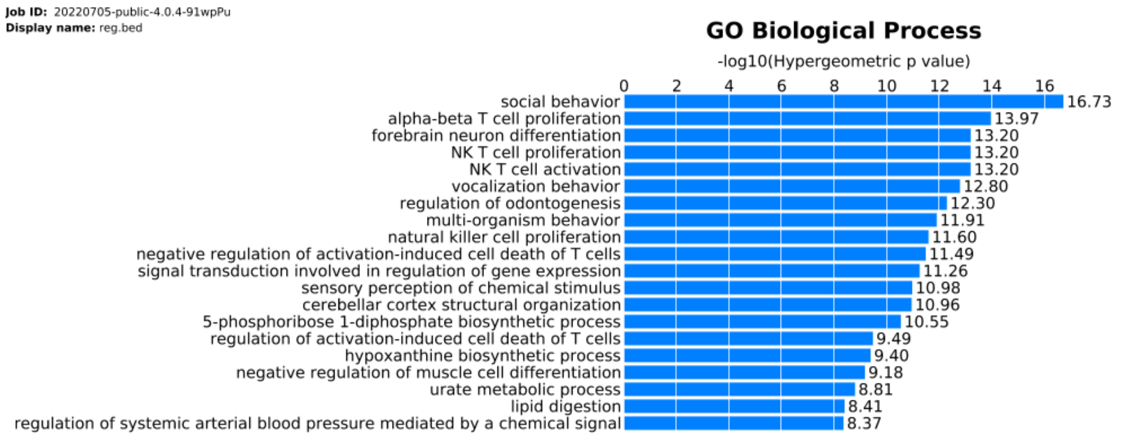

***Supplementary Figure 9: Results from simulated data.*** **A:** Accuracy of MC profiles from simulated datasets with increased coverage (y-axis: average JSD value of the MC profiles estimated between 1000 low-coverage 4-CpG datasets and the MC profile computed from the high-coverage 4-CpG dataset. x-axis: number of reads to simulate low coverage datasets. Dashed lines: gain in accuracy when increasing the coverage between 25 and 50 reads. Solid lines: interval of increased coverage to obtain the gain in accuracy observed between 25 and 50 reads). The shaded area indicates the standard deviation of the observed JSD value at a given coverage. **B:** Precision of MC profiles estimated from simulated 50 reads datasets for each 4-CpG high coverage dataset (y-axis: JSD values between MC profiles estimated from 1000 datasets' pairs; x-axis: 4-CpG datasets).

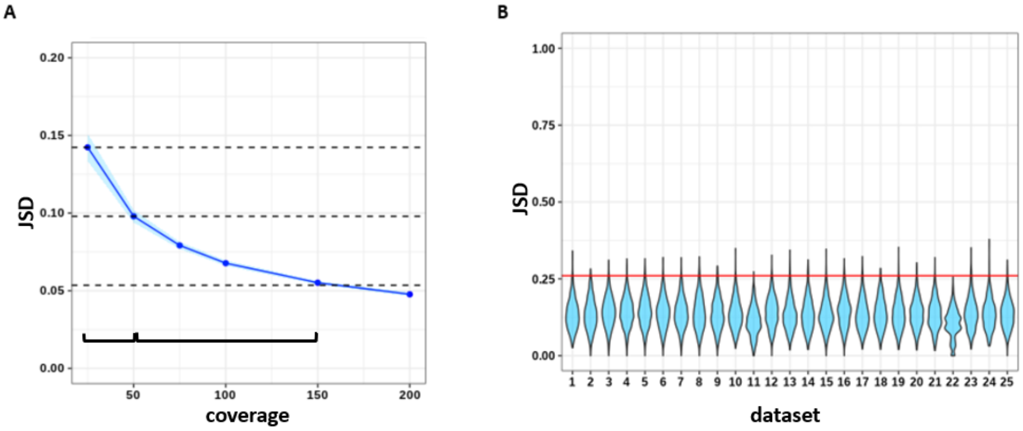

***Supplementary Figure 10: Appropriateness of data compression scheme for Dataset 1 (panel A) and Dataset 2 (panel B).*** For each MC profiles, the within class distance (WCD), i.e. the JSD between the MC profiles to the most similar Methylation Pattern (MP) is indicated on the x-axis, whereas the External Class Distance (ECD), i.e. the JSD from the second most similar MP, is shown on the y-axis. The MC profiles whose ECD/WCD ratio was higher than 1.5 (95% of MC profiles for Dataset 1 and 98% of MC profiles for Dataset 2) are colored in blue.

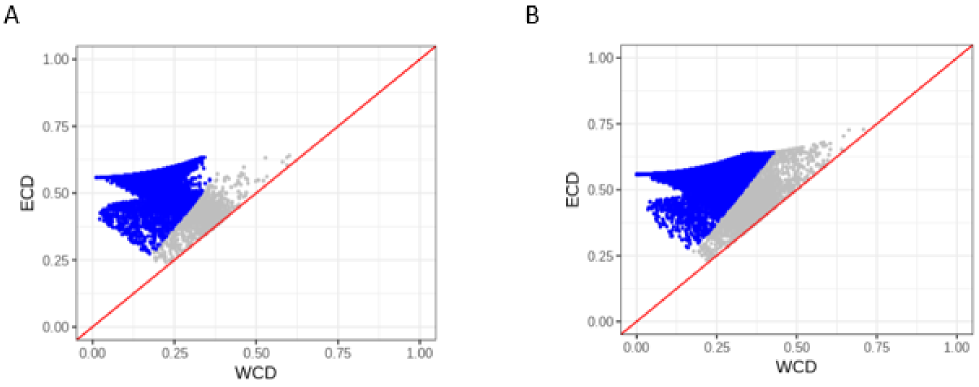

### *Supplementary Tables*

**Supplementary Table 1: RRBS datasets adopted in the study**

| **Dataset** | **GEO accession** | **Sample accessions** | **Description** |
|---|---|---|---|
| Dataset 1 | GSE130735 | GSM3752619, GSM3752620, GSM3752621 | samples from 3 WT littermate E8.5 embryos |
| Dataset 2 | GSE66121 | GSM1614765, GSM1614766, GSM1614767 | human CD19+ B-cells isolated from 3 normal controls |
| Dataset 3 | GSE53714 | GSM1299332, GSM1299333, GSM1299334 | liver samples from 3 F1 mice originating from C57BL/6J and DBA/2J strain cross |
| Dataset 4 | GSE66121 | GSM1614729, GSM1614730, GSM1614731 | human CD19+ B-cells isolated from 3 chronic lymphocytic leukemia |
| Dataset 5 | GSE72700 | GSM1868584, GSM1868589, GSM1868591 | ERRBS data from C57BL6 male mice neurons at different developmental stages (hippocampal precursors:, granule cells:; CA3 neurons:) |

*Supplementary Table 2: Description of the D-ABS data for low-coverage 4-CpG data simulation*

| Gene | Organism | Amplicon Coordinates | Genome Assembly | Primer FW | Primer RV |
|---|---|---|---|---|---|
| DAO | Human | CHR12:108879926-108880252 | GRCh38/hg38 | aaggTTtgtTTaTaggggTttgaga | ccaActcaaaAAtAcatctAccactc |
| DDOH | Human | CHR6: 110415392-110415789 | GRCh38/hg38 | aTTtaTaaatTagTtggagaaagTTTag | cctattcaAacacactcccaaactcc |
| SCRN1 | Human | CHR7:29990018-29990346 | GRCh38/hg38 | gatatggaatTttggTttagtta | ctttActaAattttatttctt |
| CDKL5 | Mouse | CHRX: 160994844-160994655 | GRCm38/mm10 | AgAgggTTAgAATAAgAAATTTTggTT | TCTAAAAACAAACTAATAAACACAC |
| DDO_R3 | Mouse | CHR10:40629085-40629513 | GRCm38/mm10 | GTttttTTaTatgtTttggagTTt | acctccctAaaaAtcatttAattcta |
| DDO_R4 | Mouse | CHR10:40629544-40629949 | GRCm38/mm10 | GtgtgtttTtgaggaggtgaTaTtTa | aActtaccctccattAAtccatAcc |
| DDO_R6 | Mouse | CHR10:40630278-40630682 | GRCm38/mm10 | TTtagtgttaaTttattagagTtgtgg | AAtacaatcccttcttAcaacaAAca |
| DDO_R7 | Mouse | CHR10:40630812-40631211 | GRCm38/mm10 | GagggagttgggTatggagTaTaTata | AactctaAAAaAcaAacacaAaAAtc |
| DLX6 | Mouse | CHR6:6864874-6865260 | GRCm38/mm10 | tTataatgTattgttagtgttggaga | tcaAtcctaaAAaaAcaAccaAttcR |
| TPH1a | Zebrafish | CHR25:8159799-8160116 | GRCz11/danRer11 | atttgTtgtTaggaggaagattaag | cacaacatcaaattctctacat |

*(*genes: human_DDO: human D-Aspartate Oxidase, mouse_DDOR4: mouse D-Aspartate Oxidase Region 4, mouse_DDOR6: mouse D-Aspartate Oxidase Region 6, mouse_DDOR7: mouse D-Aspartate Oxidase Region 7*)*

*Supplementary Table 3: Description of the simulated 4-CpGs datasets*

| gene | cpg | reads | subset | mean_met | MC_shannon | intmet |
|---|---|---|---|---|---|---|
| human_DAO | 10 | 83582 | E1 | 0.5449 | 2.2413 | 3 |
| human_DAO | 10 | 83582 | E2 | 0.6958 | 1.9836 | 4 |
| human_DAO | 10 | 71454 | E2 | 0.8345 | 1.4487 | 5 |
| human_DDO | 7 | 29333 | E1 | 0.1453 | 1.0046 | 1 |
| human_DDO | 7 | 78598 | E1 | 0.2027 | 1.3182 | 2 |
| human_SCRN1 | 14 | 49047 | E1 | 0.3857 | 1.8884 | 2 |
| human_SCRN1 | 14 | 49047 | E2 | 0.185 | 1.6757 | 1 |
| human_SCRN1 | 14 | 55839 | E1 | 0.4452 | 1.9994 | 3 |
| mouse_CDKL5 | 42 | 31845 | E1 | 0.4299 | 1.9901 | 3 |
| mouse_CDKL5 | 42 | 31845 | E2 | 0.3033 | 2.0264 | 2 |
| mouse_DDOR3 | 9 | 181424 | E1 | 0.9207 | 1.0398 | 5 |
| mouse_DDOR3 | 9 | 181424 | E2 | 0.6964 | 2.0182 | 4 |
| mouse_DDOR4 | 6 | 117541 | E1 | 0.6354 | 2.2042 | 4 |
| mouse_DDOR4 | 6 | 267331 | E1 | 0.5302 | 2.2878 | 3 |
| mouse_DDOR4 | 6 | 87085 | E1 | 0.3869 | 2.1063 | 2 |
| mouse_DDOR4 | 6 | 101393 | E1 | 0.1893 | 1.7031 | 1 |
| mouse_DDOR6 | 6 | 127610 | E1 | 0.5561 | 2.2882 | 3 |
| mouse_DDOR6 | 6 | 26696 | E1 | 0.6517 | 2.1715 | 4 |
| mouse_DDOR6 | 6 | 86888 | E1 | 0.3216 | 2.1247 | 2 |
| mouse_DDOR7 | 9 | 94045 | E1 | 0.6973 | 2.0404 | 4 |
| mouse_DDOR7 | 9 | 217679 | E2 | 0.8002 | 1.7244 | 5 |
| mouse_DLX6 | 16 | 115382 | E1 | 0.0248 | 0.4587 | 1 |
| mouse_DLX6 | 16 | 90607 | E4 | 0.2419 | 1.8148 | 2 |
| zebra_TPH1a | 11 | 39642 | E1 | 0.5734 | 2.2836 | 3 |
| zebra_TPH1a | 11 | 56543 | E1 | 0.6467 | 2.1782 | 4 |

(genes: human_DDO: human D-Aspartate Oxidase, mouse_DDOR4: mouse D-Aspartate Oxidase Region 4, mouse_DDOR6: mouse D-Aspartate Oxidase Region 6, mouse_DDOR7: mouse D-Aspartate Oxidase Region 7. column names: subset: simulated short-read region made up of 4 CpG sites, mean_met: average methylation of the simulated high-coverage short-read dataset, MC_shannon: shannon entropy computed on the high-coverage short-read dataset, intmet: average methylation-based group.)

***Supplementary Table 4: simulated datasets based grouped by average methylation***

| group | number of datasets | average methylation | average coverage |
|---|---|---|---|
| G1 | 4 | 0-0.2 | 100974.8 |
| G2 | 6 | 0.2-0.4 | 93305.17 |
| G3 | 6 | 0.4-0.6 | 156852.3 |
| G4 | 6 | 0.6-0.8 | 73788.75 |
| G5 | 3 | 0.8-1 | 70678.33 |
